# Cannabidiol Reduces D1 and D2 Medium Spiny Neuron Excitability in the Nucleus Accumbens Core

**DOI:** 10.64898/2025.12.02.691956

**Authors:** Zev E. Jarrett, Brad A. Grueter

**Author notes:** **Correspondence to:** Brad A. Grueter, Ph.D., Department of Anesthesiology, 2213 Garland Avenue, P435H MRB IV, Vanderbilt University Medical Center, Nashville, TN 37232-0413.

## Abstract

Cannabidiol (CBD) is a non-psychotomimetic phytocannabinoid constituent of the cannabis plant that shows promise for the treatment of a variety of neuropsychiatric disorders such as anxiety disorders, post-traumatic stress disorder, and substance use disorders. The nucleus accumbens (NAc) is a key brain region in the etiologies of these disorders and is actively modulated by CBD. Prior research has established that CBD alters the molecular composition of the NAc, but none have assessed how CBD affects NAc neuronal function. In this study, we demonstrate that CBD significantly decreases D1 and D2 medium spiny neuron membrane excitability, broadens action potentials, and has no effect on spontaneous excitatory synaptic transmission in the NAc core. These data enhance our understanding of CBD’s physiological effects and provide mechanistic insight into its therapeutic potential.

## Introduction

Cannabidiol (CBD) is a major non-psychotomimetic phytocannabinoid constituent of the cannabis plant that has seen significant increases in commercialization and recreational use. In 2022, 20% of adults in the U.S. reported using CBD/hemp-derived products^1^. However, there is a paucity of knowledge of the mechanisms behind CBD’s putative therapeutic effects. CBD has been FDA approved for the treatment of certain pediatric-onset epilepsies and shows promise for the treatment of a variety of neuropsychiatric disorders such as anxiety disorders, post-traumatic stress disorder, and substance use disorders^2–4^. The nucleus accumbens (NAc) is a key brain region in the etiologies of these disorders and is actively modulated by CBD ^5–11^. For example, CBD decreases alcohol craving and cue-induced NAc activation in human subjects with alcohol use disorder^12^. Preclinical work has correlated improvements in constructs of neuropsychiatric disorders with changes in the molecular composition of the NAc^13–17^. However, there is a gap in knowledge of the functional consequences of CBD on NAc neuronal properties.

95% of NAc neurons are medium spiny projection neurons (MSNs). MSNs can be subdivided into two types: dopamine receptor type 1 and type 2-expressing (D1 and D2 MSNs). NAc D1 and D2 MSNs serve as a key node in the integration of information from a variety of brain regions involved in neuropsychiatric disorders such as the hippocampus, amygdala, prefrontal cortex, and thalamus^10^. Therefore, assessing how CBD affects MSN function may elucidate mechanisms by which CBD exerts its therapeutic effects. In this study, we utilized whole-cell patch-clamp electrophysiology in acute brain slices from transgenic mice to assess the functional effects of CBD on D1 and D2 MSNs within the NAc core (NAcC). We found that CBD significantly decreases D1 and D2 MSN membrane excitability, broadens action potentials, increases afterhyperpolarization, and has no effect on spontaneous excitatory synaptic transmission. These results enhance our understanding of CBD’s effects on NAc physiology and provide mechanistic insight into its therapeutic potential.

## Methods

### Subjects

Experiments were performed adhering to the National Institutes of Health guidelines for the Care and Use of Laboratory Animals and were approved by the Vanderbilt University Institutional Animal Care and Use Committee. All experiments used male and female D1^tdtomato^ (Jackson Laboratories, Cat. #016204) and/or D2^GFP^ mice (Tg(Drd2-EGPF)S118Gsat) aged 8-12 weeks.

### Ex Vivo Electrophysiology

To collect acute ex vivo brain slices containing NAcC, mice were anesthetized with isoflurane and rapidly decapitated. Dissected tissue was placed into ice-cold oxygenated dissecting solution and sectioned at 250 μM for sagittal slices of NAcC using the Leica VT1200S Vibratome. The dissecting solution consisted of (in mM): 93 N-methyl-D-glucamine (NMDG), 2.5 KCl, 20 HEPES, 10 MgCl_2_, 1.2 NaH_2_PO_4_·H_2_O, 30 NaHCO_3_, 0.5 CaCl_2_· 2H_2_0, 25 glucose, 3 Na-pyruvate, 5 ascorbic acid. The aCSF consisted of (in mM): 118.9 NaCl, 2.5 KCl, 1.3 MgCl_2_·6H_2_O, 2.5 CaCl_2_, 1 NaH_2_PO_4_, 26.2 NaHCO_3_, 11 glucose. Cannabidiol (Cayman Chemical, Cat. #90080) was dissolved in DMSO at a stock concentration of 20 mM. For incubation experiments, slices were incubated for a minimum of 1 hr with aCSF containing either 20 µM CBD or equivalent dimethyl sulfoxide (DMSO). For acute application experiments, baseline was obtained with aCSF prior to 10 min perfusion of 20 µM CBD or DMSO. Slices were acclimated to the rig for 30 min prior to recording, perfused at a flow rate of 2-3 mL/min, and maintained at 29-31 °C.

Whole-cell patch clamp electrophysiology was performed using a CV-7B headstage, Multiclamp 700B Amplifier, and Axopatch Digidata 1550 digitizer. Cells were visualized at 40X magnification using an upright microscope (Scientifica) that allowed for both infrared-differential interference contrast and fluorescence optics. D1 MSNs were identified as fluorescent cells at 550nM in D1^tdtomato^ mice and non-fluorescent cells at 470nM in D2^GFP^ mice. D2 MSNs were identified as fluorescent cells at 470nM in D2^GFP^ mice and non-fluorescent cells at 550nM in D1^tdtomato^ mice. 3-5 MΩ glass recording micropipettes (Sutter P1000 Micropipette Puller) filled with potassium gluconate internal solution were used for patching. The internal solution consisted of (in mM): 135 K-gluconate, 5 NaCl, 2 MgCl_2_, 10 HEPES, 0.2 EGTA, 2.5 Mg_2_-ATP, 0.2 Na_2_-GTP. All cells were voltage clamped at -70 mV for 5 min of dialysis prior to recording. Current clamp experiments were performed by clamping cells at a baseline holding current to rest at -70 mV. The excitability protocol consisted of 800 ms square wave current every 5 sec from -50 pA to 400 pA in 25 pA increments. Voltage clamp experiments were performed by voltage clamping cells at -70 mV for 5 min of gap free recording. Cells were excluded from analysis if they had a baseline series resistance ≥ 25 MOhm or change in series resistance ≥ 20%.

### Quantification and Statistical Analysis

Electrophysiology data were initially processed in Clampfit 10 software (Molecular Devices) and quantified in Microsoft Excel. Data were analyzed and graphs created with GraphPad Prism 10 software. The significance threshold for all analyses was *p* < 0.05. Data are shown as mean ± SEM.

## Results

### Cannabidiol incubation reduces NAcC D1 and D2 MSN membrane excitability

CBD interacts with various substrates that can influence membrane excitability and has been shown to inhibit action potential generation in other brain regions^18^. To determine if CBD affects MSN excitability in the NAcC, we incubated acute brain slices containing NAcC in 20 µM CBD or vehicle solution (aCSF with proportional DMSO) for a minimum of 1 hr and recorded action potential frequency in response to square wave current injection. Current injections for each cell ranged from -50 to 400 pA in 25 pA increments. D1 and D2 MSNs incubated in CBD showed significantly decreased action potential firing frequency compared to vehicle control (Figures 1, 2 A). Next, we assessed action potential properties in response to CBD. We analyzed the half-width, maximum decay slope, afterhyperpolarization peak, and maximum rise slope of the first action potential generated under 200 pA current injection. For D1 and D2 MSNs, 20 µM CBD significantly increased the half-width, decreased the maximum decay slope, decreased the afterhyperpolarization peak, and had no effect on maximum rise slope (Figures 1, 2 B-E). These results implicate potassium channels as a CBD target, as half-width, afterhyperpolarization peak, and maximum decay slope are indicators of potassium channel activity.

**Figure 1.**
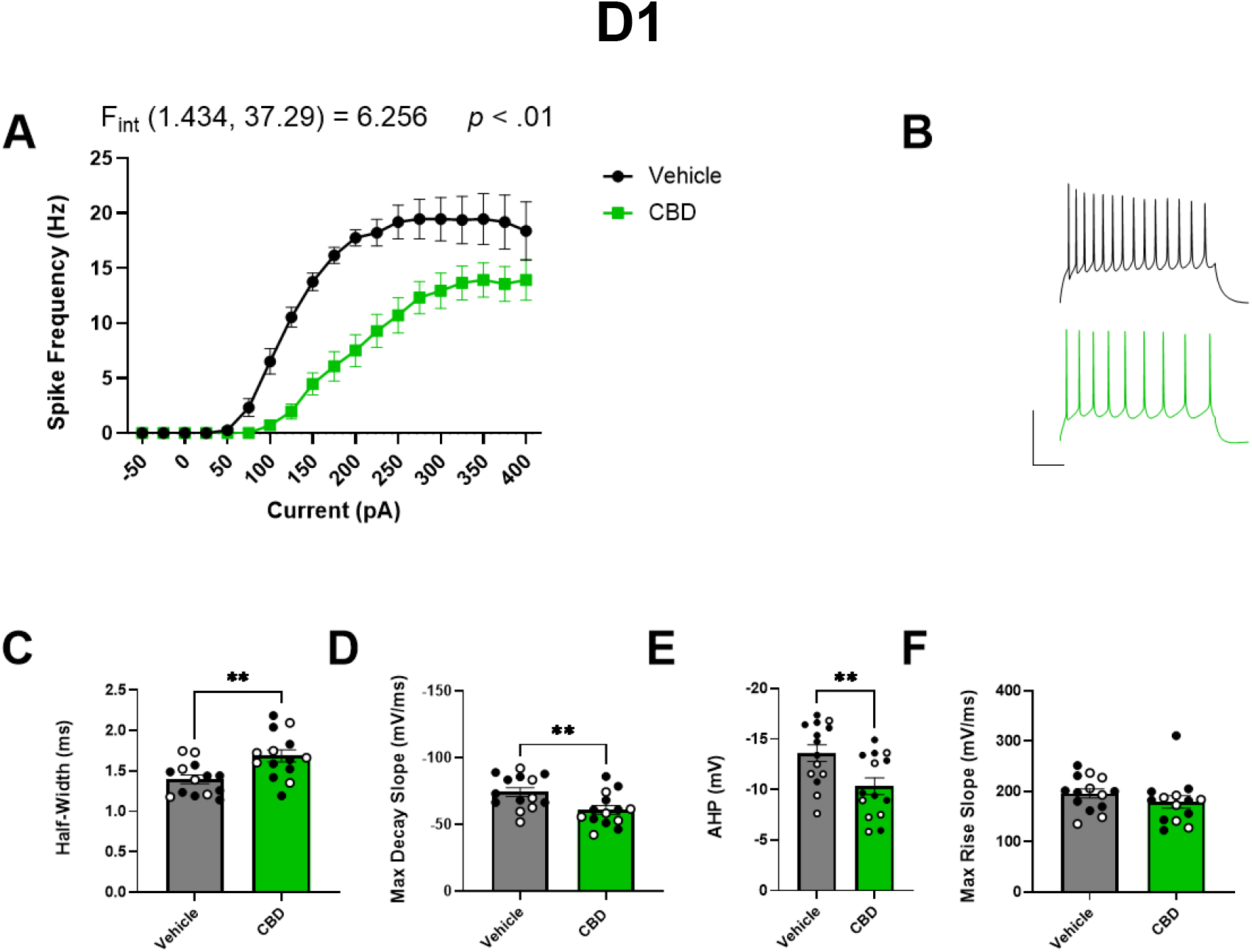
20 uM CBD incubation decreases NAcC D1 MSN excitability. (A) Effect of 20 µM CBD incubation on D1 MSN excitability. (B) Representative traces of D1 MSNs incubated in either DMSO or CBD (scale = 100 ms, 50 mV). For C-F, the first action potential generated at 200 pA is analyzed. (C-F) Effect of 20 µM CBD incubation on D1 MSN (C) half-width, (D) maximum decay slope, (E) afterhyperpolarization peak, (F) maximum rise slope. Black and white points denote male and female mice, respectively. Data analyzed by (A) two-way repeated measures ANOVA or (C-F) independent two-tailed *t*-test (\*\**p* < .01).

**Figure 2.**
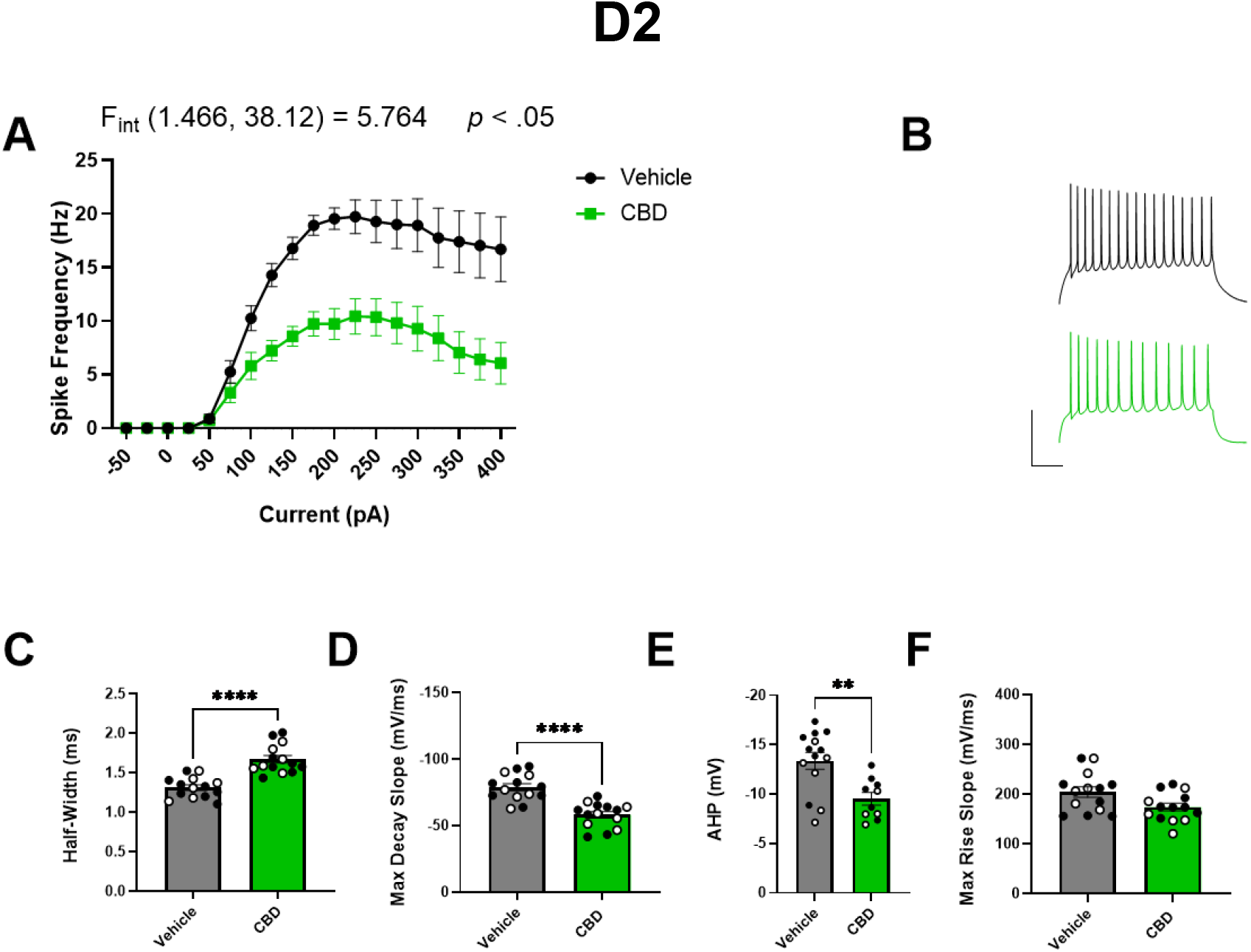
20 uM CBD incubation decreases NAcC D2 MSN excitability. (A) Effect of 20 µM CBD incubation on D2 MSN excitability. (B) Representative traces of D2 MSNs incubated in either DMSO or CBD (scale = 100 ms, 50 mV). For C-F, the first action potential generated at 200 pA is analyzed. (C-F) Effect of 20 µM CBD incubation on D2 MSN (C) half-width, (D) maximum decay slope, (E) afterhyperpolarization peak, (F) maximum rise slope. Black and white points denote male and female mice, respectively. Data analyzed by (A) two-way repeated measures ANOVA or (C-F) independent two-tailed *t*-test (\*\**p* < .01, \*\*\*\**p* < .0001).

### Acute cannabidiol exposure decreases NAcC D1 and D2 MSN membrane excitability at high levels of current

To evaluate the time course of CBD’s effects on MSN excitability, we compared action potential frequency pre and post 10 min 20 µM CBD or DMSO bath application. D1 and D2 MSNs showed a significant decrease in frequency at high levels of current injection with CBD but not DMSO (Figures 3, 4 A-C). D1 MSNs displayed no change in half-width or maximum decay slope with CBD or DMSO, a significant decrease in afterhyperpolarization peak with CBD but not DMSO, and a significant decrease in maximum rise slope with both CBD and DMSO (Figure 3, D-G, K-N). D2 MSNs displayed no change in half-width with CBD or DMSO, a significant decrease in maximum decay slope with CBD but not DMSO, a significant decrease in afterhyperpolarization peak with both CBD and DMSO, and no change in maximum rise slope with CBD or DMSO (Figure 4, D-G, K-N). These results suggest that CBD may have low potency at its target or activates a signaling cascade that regulates ion channel function on a slower time scale.

**Figure 3.**
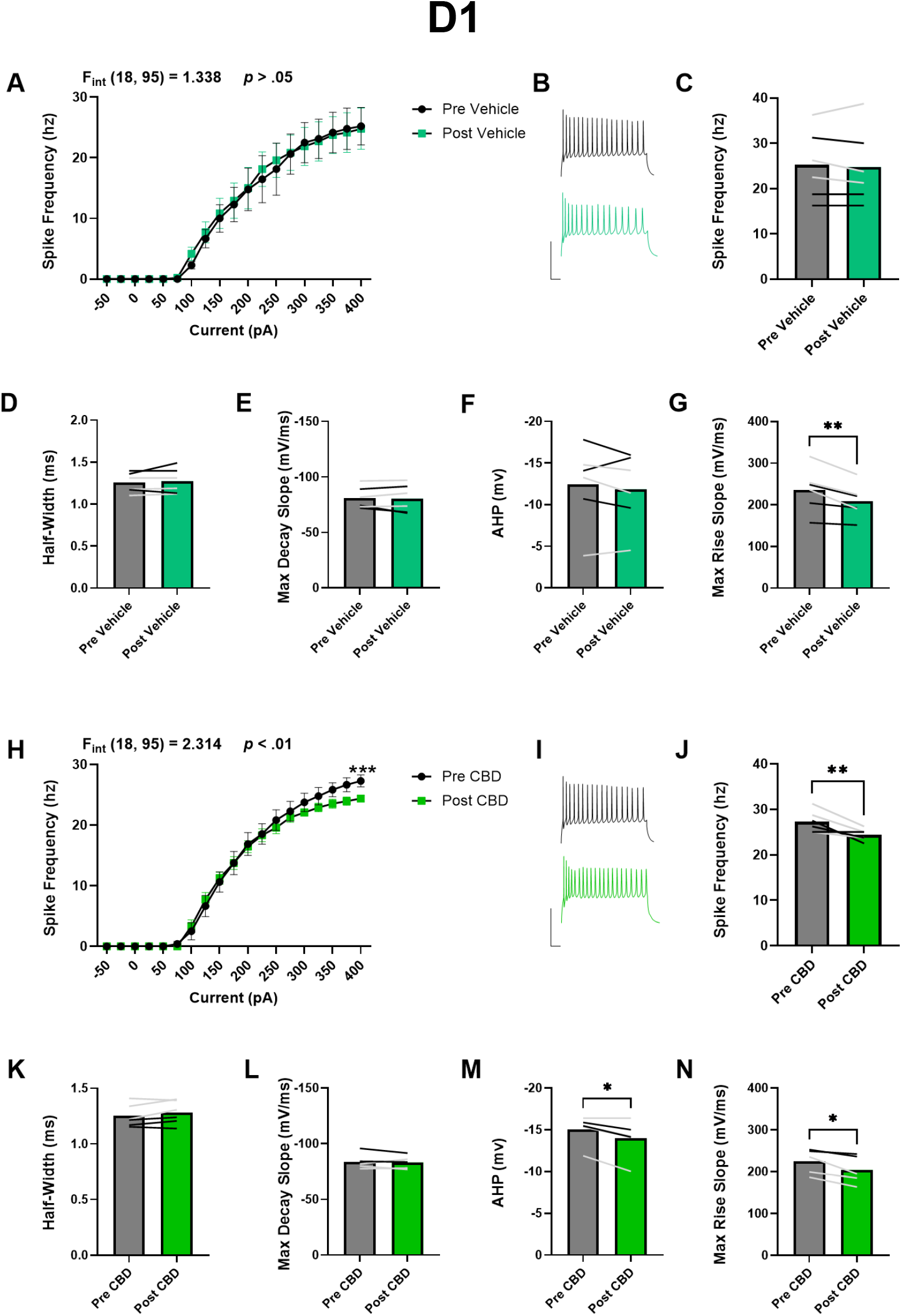
Acute 20 uM CBD decreases NAcC D1 MSN excitability at high levels of current. Acute 10 min application of 20 µM CBD or equivalent DMSO (A) Effect of acute DMSO on D1 MSN excitability. (B) Representative traces of D1 MSNs pre and post DMSO (scale = 100 ms, 50 mV). (C) Action potential frequency at 400pA current injection pre and post DMSO. (D-G) Effect of acute DMSO on D1 MSN (D) half-width, (E) maximum decay slope, (F) afterhyperpolarization peak, (G) maximum rise slope. (H-N) same as (A-G) for CBD. Black and white lines denote male and female mice, respectively. Data analyzed by (A, H) two-way repeated measures ANOVA with Holm-Šídák multiple comparisons, (C, J) Holm-Šídák multiple comparison from (A, H), (D-G, K-N) paired two-tailed *t*-test (\**p* < .05, \*\**p* < .01).

**Figure 4.**
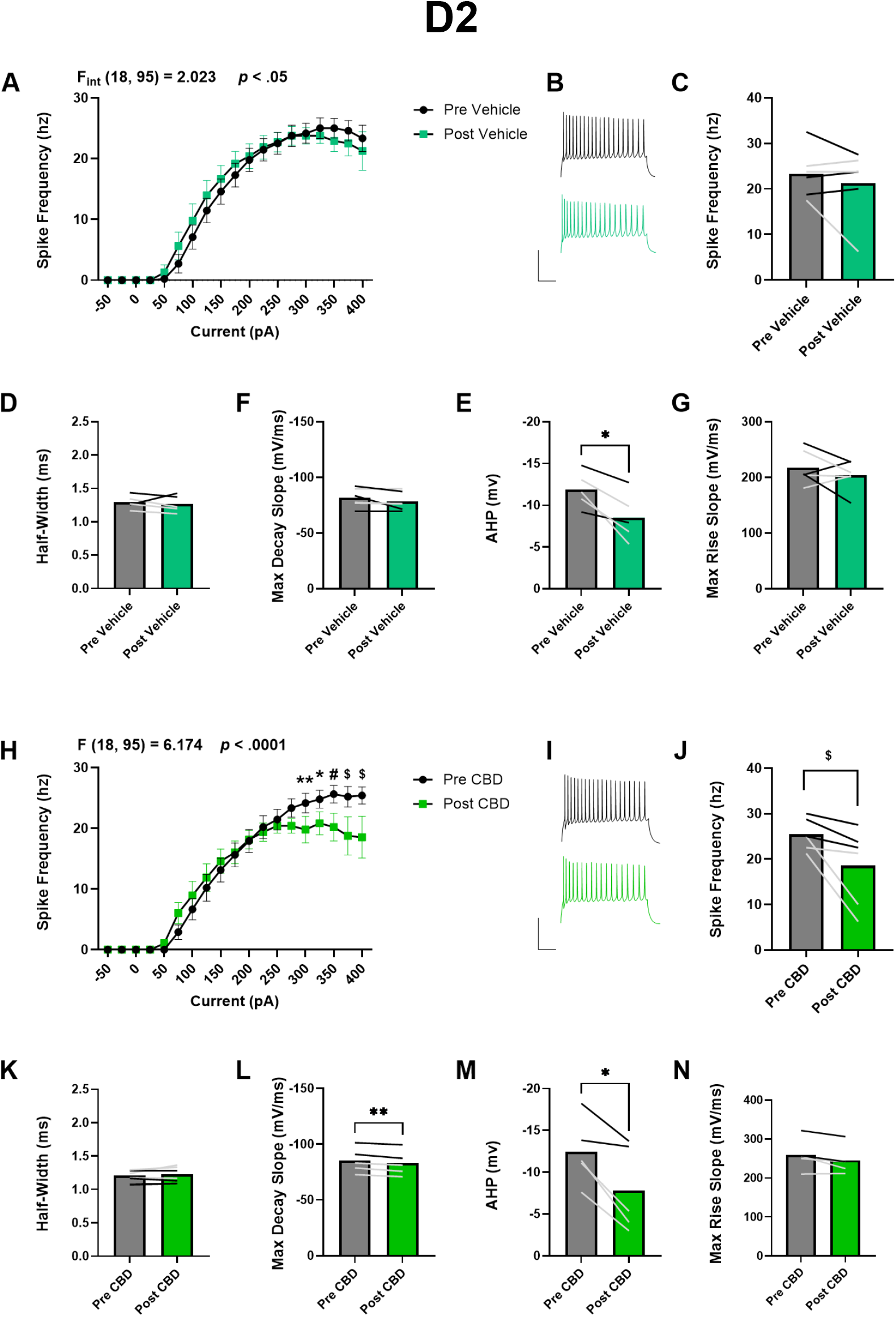
Acute 20 uM CBD decreases NAcC D2 MSN excitability at high levels of current. Acute 10 min application of 20 µM CBD or equivalent DMSO (A) Effect of acute DMSO on D2 MSN excitability. (B) Representative traces of D1 MSNs pre and post DMSO (scale = 100 ms, 50 mV). (C) Action potential frequency at 400pA current injection pre and post DMSO. (D-G) Effect of acute DMSO on D1 MSN (D) half-width, (E) maximum decay slope, (F) afterhyperpolarization peak, (G) maximum rise slope. (H-N) same as (A-G) for CBD. Black and white lines denote male and female mice, respectively. Data analyzed by (A, H) two-way repeated measures ANOVA with Holm-Šídák multiple comparisons, (C, J) Holm-Šídák multiple comparison from (A, H), (D-G, K-N) paired two-tailed *t*-test (\**p* < .05, \*\**p* < .01, #*p* < .001, $*p* < .0001).

### Cannabidiol incubation has no effect on NAcC D1 and D2 MSN spontaneous excitatory neurotransmission

Prior research has shown that CBD can alter spontaneous excitatory post synaptic currents (sEPSCs) in multiple brain regions^18–20^. To determine if CBD affects sEPSCs in the NAcC, we incubated acute brain slices containing NAcC in 20 µM CBD or DMSO and recorded sEPSCs. We found no significant difference in sEPSC frequency or amplitude for D1 and D2 MSNs, suggesting that 20 µM CBD does not affect glutamatergic presynaptic release probability or postsynaptic AMPA receptor function in NAcC MSNs (Figure 5).

**Figure 5.**
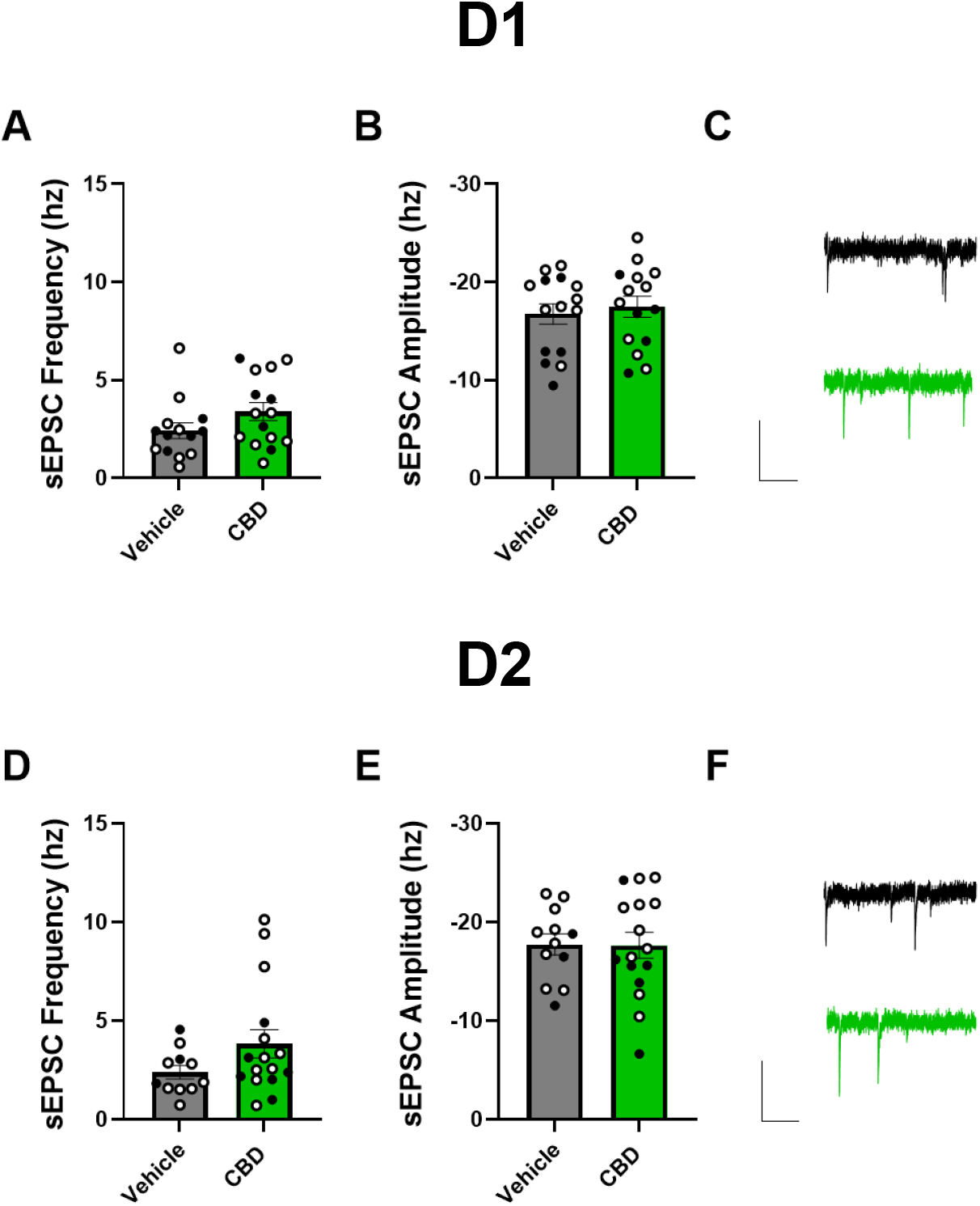
20 uM CBD incubation has no effect on NAcC D1 or D2 MSN spontaneous excitatory transmission. (A) Effect of 20 µM CBD or DMSO on sEPSC frequency in D1 MSNs. (B) Effect of 20 µM CBD or DMSO on sEPSC amplitude in D1 MSNs. (C) Representative traces of CBD and DMSO-incubated D1 MSN sEPSCs (scale = 250 ms, 20 pA). (D-F) Same as A-C but for D2 MSNs. Black and white points denote male and female mice, respectively. Data analyzed by independent two-tailed *t*-test.

## Discussion

This study is the first to investigate the consequences of CBD exposure on electrophysiological properties of NAcC MSNs. We establish that CBD reduces NAcC D1 and D2 MSN membrane excitability, broadens action potentials, and decreases afterhyperpolarization while having no detectable effects on spontaneous excitatory synaptic transmission. Because CBD incubation resulted in a robust decrease in excitability while acute application only showed effects at high levels of current injection, CBD may have low potency at its target or activate a signaling cascade that regulates ion channel function on a slower time scale (Figures 1-4).

Our finding that CBD causes reduced membrane excitability in D1 and D2 MSNs (Figures 1, 2) is consistent with results from striatal cultures^21^. Additionally, CBD has been shown to reduce excitability of neurons in the central and basolateral amygdala, hippocampus, dentate gyrus, and dorsal root ganglia^18,20,22–24^. The molecular mechanism(s) underlying CBD’s inhibition of neuronal excitability are unclear. One potential site of action is voltage gated sodium channel 1.6 (NaV1.6). NaV1.6 regulates MSN excitability in the NAc, and CBD inhibits NaV1.6 resurgent and persistent current in striatal cultures^21,25,26^. While our analysis found no effect of CBD on maximum rise slope, this metric is an indicator of total sodium current during the depolarization phase and is thus not able to detect a change in resurgent or persistent current. Indeed, CBD has no effect on NaV1.6 peak transient current^21^. Our action potential analysis suggests that CBD targets potassium channels in the NAcC, as half-width, maximum decay slope, and afterhyperpolarization peak are potassium channel dependent (Figures 1, 2). In vitro data suggests multiple potassium channels as CBD targets. For example, Kv7.2/7.3-mediated low-voltage activated M-current is enhanced by CBD in vitro, Kv7.2/7.3 enhancement decreases excitability, and these channels are expressed on NAc MSNs^27–32^. Thus, CBD-enhanced M-current may be responsible for our results. However, our finding of increased half-width and decreased maximum decay slope and afterhyperpolarization peak would suggest inhibited M-current (Figures 1, 2). Still, other potassium channels may mediate these action potential properties while CBD decreases MSN excitability by enhancing Kv7.2/7.3. Additional potassium channels expressed on NAc MSNs that are targeted by CBD include Trek-1 and BK channels^33–38^. CBD inhibits both mechanosensitive Trek-1 and calcium-activated BK channels in vitro and their respective knockouts lead to increased half-width^33,36,39,40^. However, Trek-1 and BK channel activation has been shown to decrease neuronal excitability in other brain regions^40,41^. Thus, one would predict increased excitability if CBD were inhibiting these channels in MSNs. Another ion channel targeted by CBD that is expressed by NAc MSNs is the non-selective cation channel transient receptor potential vanilloid 1 (TRPV1)^42^. CBD is a TRPV1 agonist and altered conductance by TRPV1 activation may contribute to the phenotypes we uncovered^42–44^. We have discussed a myriad of previously identified membrane effector systems linked to CBD action. However, it is possible that a unique signaling mechanism is recruited by CBD in NAcC MSNs.

Acute 10 min CBD application led to a similar decrease in NAcC D1 and D2 MSN excitability as incubation, but only at high levels of current injection. Most action potential properties were unaffected by acute CBD (either noeffect with CBD or DMSO or a change in both). However, D1 MSNs showed a significant decrease in afterhyperpolarization peak and D2 MSNs showed a significant decrease in maximum decay slope with acute CBD but not DMSO. These results are in the same direction as with incubation. However, most action potential properties were still unaffected by acute CBD and the two that showed effects were distinct between D1 and D2 MSNs (Figures 4, 5). We believe that the dichotomy between incubation and acute exposure in terms of excitability and action potential properties suggests that distinct mechanisms may be responsible for lowering excitability and altering action potential kinetics. Conversely, action potential broadening/increased afterhyperpolarization may be a consequence of reducing excitability that only emerges with longer incubation. The fact that CBD requires incubation to reach its full effects suggests that it may have low potency at its target (e.g. sodium/potassium channels) or work on a longer time scale through second messenger pathways. Beyond ion channels, CBD can act as an: inhibitor of the reuptake/metabolism of anandamide (cannabinoid receptor 1 [CB1] endogenous agonist), CB1 negative allosteric modulator, G-protein-coupled receptor 55 (GPR55) antagonist, 5-HT_1A_ agonist, peroxisome proliferator-activated receptor (PPAR) gamma agonist, mu and delta opioid receptor negative allosteric modulator, adenosine 1 receptor (A1) agonist and reuptake inhibitor, and more^45–47^. With such a wide array of targets, some expressed on NAc MSNs (e.g. GPR55, A1) and some not (e.g. 5-HT_1A_, PPAR), there is a high likelihood of complex synergistic effects of multiple receptor systems to lead to our results. Suggesting the complexity of CBD-mediated altered excitability, CBD has an inhibitory effect on action potential firing frequency in almost every cell type in which it has been assessed except parvalbumin-expressing fast-spiking interneurons, in which it conversely leads to increased membrane excitability^18,20,22,23^.

We found that CBD caused no change in spontaneous excitatory synaptic transmission in D1 and D2 MSNs in the NAcC. Prior literature has established that CBD affects synaptic transmission in a brain region and cell-type specific manner^18,20,23^. Our work suggests that CBD’s effect on NAcC MSNs may be constrained to membrane excitability with no effect on synaptic transmission. However, we only assessed sEPSCs. There are cell types such as somatostatin neurons in the central amygdala that show a reduction in spontaneous inhibitory postsynaptic currents (sIPSCs) but no change in sEPSCs with CBD incubation^18^. Future work should assess the effects of CBD on sIPSCs in NAcC MSNs. Additionally, there is the possibility of CBD acting as a depotentiator of synaptic plasticity. CBD has been shown to normalize increases in sEPSC and decreases in sIPSC frequency caused by the endogenous membrane phospholipid lysophosphatidylinositol in CA1 pyramidal neurons with no effect by itself^19^. Therefore, CBD may be able to normalize plastic changes in spontaneous synaptic transmission in NAc MSNs as can be caused by chronic cocaine or morphine exposure^48,49^.

The ability of CBD to decrease NAc MSN excitability shows therapeutic promise for the treatment of a variety of disease states. For example, increased NAc MSN (core and/or shell) excitability has been found in response to exposure/withdrawal from drugs of abuse, acute restraint stress, juvenile social isolation, fear conditioning, and obesity^50–56^. Indeed, CBD has shown promise as a treatment for anxiety disorders, post-traumatic stress disorder, and substance use disorders^2,3^. Clinically, CBD has been shown to reduce alcohol craving and decrease functional NAc activity in response to alcohol-paired cues in subjects with alcohol use disorder^12^. Our finding of decreased NAcC MSN excitability provides a physiological mechanism for decreased alcohol-paired cue-induced NAc activation.

In conclusion, this work is the first to investigate the functional physiological effects of CBD in the NAc. We establish that CBD massively inhibits D1 and D2 MSN membrane excitability in the NAcC and alters potassium channel-dependent action potential kinetics. Future work should further evaluate the molecular mechanisms underlying these effects and assess whether inhibition of NAc excitability by CBD can improve preclinical disease models.

## Author contributions

Conceptualization (ZEJ, BAG), Data curation (ZEJ), Funding acquisition (BAG), Methodology (ZEJ), Writing – original draft (ZEJ), Writing – review and editing (BAG).

## Funding Sources

This work was supported by NIDA R01DA040630 (BAG) NIMH T32 GM149363 (ZEJ)

## Conflict of Interest

Authors report no conflict of interest

## Notes

### Competing Interest Statement

The authors have declared no competing interest.

## References

1. Choi NG, Marti CN, Choi BY. Prevalence of cannabidiol use and correlates in U.S. adults. Drug and alcohol dependence reports. 2024 Dec;13:100289. PMID: 39831101

2. Navarrete F, García-Gutiérrez MS, Gasparyan A, Austrich-Olivares A, Manzanares J. Role of Cannabidiol in the Therapeutic Intervention for Substance Use Disorders. Front Pharmacol. 2021;12:626010. PMID: 34093179

3. Han K, Wang JY, Wang PY, Peng YCH. Therapeutic potential of cannabidiol (CBD) in anxiety disorders: A systematic review and meta-analysis. Psychiatry Res. 2024 Sep;339:116049. PMID: 38924898

4. Silva GD, Del Guerra FB, de Oliveira Lelis M, Pinto LF. Cannabidiol in the Treatment of Epilepsy: A Focused Review of Evidence and Gaps. Front Neurol. 2020;11:531939. PMID: 33192966

5. Sturm V, Lenartz D, Koulousakis A, Treuer H, Herholz K, Klein JC, Klosterkötter J. The nucleusaccumbens: a target for deep brain stimulation in obsessive–compulsive- and anxiety-disorders. J Chem Neuroanat. 2003 Dec;26(4):293–299.

6. Burkhouse KL, Jagan Jimmy, Defelice N, Klumpp H, Ajilore O, Hosseini B, Fitzgerald KD, Monk CS, Phan KL. Nucleus accumbens volume as a predictor of anxiety symptom improvement following CBT and SSRI treatment in two independent samples. Neuropsychopharmacology. 2020 Feb;45(3):561–569. PMID: 31756730

7. Chen Y, Yan P, Wei S, Zhu Y, Lai J, Zhou Q. Ketamine metabolite alleviates morphine withdrawal-induced anxiety via modulating nucleus accumbens parvalbumin neurons in male mice. Neurobiol Dis. 2023 Oct 1;186:106279. PMID: 37661023

8. Sailer U, Robinson S, Fischmeister FPS, König D, Oppenauer C, Lueger-Schuster B, Moser E, Kryspin-Exner I, Bauer H. Altered reward processing in the nucleus accumbens and mesial prefrontal cortex of patients with posttraumatic stress disorder. Neuropsychologia. 2008 Sep;46(11):2836–44. PMID: 18597797

9. Yu YH, Lim YS, Ou CY, Chang KC, Tsai AC, Chang FC, Huang ACW. The Medial Prefrontal Cortex, Nucleus Accumbens, Basolateral Amygdala, and Hippocampus Regulate the Amelioration of Environmental Enrichment and Cue in Fear Behavior in the Animal Model of PTSD. Behavioural neurology. 2022;2022:7331714. PMID: 35178125

10. Turner BD, Kashima DT, Manz KM, Grueter CA, Grueter BA. Synaptic Plasticity in the Nucleus Accumbens: Lessons Learned from Experience. ACS Chem Neurosci. 2018 Sep 19;9(9):2114– 2126. PMID: 29280617

11. Volkow ND, Michaelides M, Baler R. The Neuroscience of Drug Reward and Addiction. Physiol Rev. 2019 Oct 1;99(4):2115–2140. PMID: 31507244

12. Zimmermann S, Teetzmann A, Baeßler J, Schreckenberger L, Zaiser J, Pfisterer M, Stenger M, Bach P. Acute cannabidiol administration reduces alcohol craving and cue-induced nucleus accumbens activation in individuals with alcohol use disorder: the double-blind randomized controlled ICONIC trial. Mol Psychiatry. 2025 Jun;30(6):2612–2619. PMID: 39668256

13. Liu L, Wang C, Wang H, Miao L, Xie T, Tian Y, Li X, Huang Y, Zeng X, Zhu B. Identification of the circRNA-miRNA-mRNA network for treating methamphetamine-induced relapse and behavioral sensitization with cannabidiol. CNS Neurosci Ther. 2024 May;30(5):e14737. PMID: 38702929

14. Ferland JMN, Chisholm A, Abdalla J, Cinar R, Johnson C, Bradshaw HB, Hurd YL. Cannabidiol abrogates cue-induced anxiety associated with normalization of mitochondria-specific transcripts and linoleic acid in the nucleus accumbens shell. Mol Psychiatry. 2025 Jun;30(6):2718–2728. PMID: 39815058

15. Navarrete F, Gasparyan A, Manzanares J. CBD-mediated regulation of heroin withdrawal-induced behavioural and molecular changes in mice. Addiction biology. 2022 Mar;27(2):e13150. PMID: 35229949

16. Dirik S, Doyle MR, Wood CP, Campo P, Martinez AR, Fannon M, Balaguer MG, Seely S, Montoya BA, Cook GMR, Palermo GM, Lin J, Sist MD, Naghshineh PK, Lan Z, Rahman SRMU, Suhandynata R, Schweitzer P, Kallupi M, de Guglielmo G. Cannabidiol mitigates alcohol dependence and withdrawal with neuroprotective effects in the basolateral amygdala and striatum.Neuropsychopharmacology. 2025 Jul 10; PMID: 40640509

17. Chisholm A, Ferland JMN, Ellis RJ, Hurd YL. Cannabidiol attenuates heroin seeking in male rats associated with normalization of discrete neurobiological signatures within the nucleusaccumbens with subregional specificity. Biol Psychiatry. 2025 Sep 22; PMID: 40992584

18. Winters ND, Yasmin F, Kondev V, Grueter BA, Patel S. Cannabidiol Differentially Modulates Synaptic Release and Cellular Excitability in Amygdala Subnuclei. ACS Chem Neurosci. 2023 Jun 7;14(11):2008–2015.

19. Rosenberg EC, Chamberland S, Bazelot M, Nebet ER, Wang X, McKenzie S, Jain S, Greenhill S, Wilson M, Marley N, Salah A, Bailey S, Patra PH, Rose R, Chenouard N, Sun SED, Jones D, Buzsáki G, Devinsky O, Woodhall G, Scharfman HE, Whalley BJ, Tsien RW. Cannabidiol modulates excitatory-inhibitory ratio to counter hippocampal hyperactivity. Neuron. 2023 Apr 19;111(8):1282-1300.e8. PMID: 36787750

20. Kaplan JS, Stella N, Catterall WA, Westenbroek RE. Cannabidiol attenuates seizures and social deficits in a mouse model of Dravet syndrome. Proc Natl Acad Sci U S A. 2017 Oct 17;114(42):11229–11234. PMID: 28973916

21. Patel RR, Barbosa C, Brustovetsky T, Brustovetsky N, Cummins TR. Aberrant epilepsy-associated mutant Nav1.6 sodium channel activity can be targeted with cannabidiol. Brain. 2016 Aug;139(Pt 8):2164–81. PMID: 27267376

22. Chamberland S, Rosenberg EC, Nebet ER, Devinsky O, Tsien RW. Cannabidiol dampens propagation of hippocampal hyperactivity and differentially modulates feedforward and feedback inhibition. bioRxiv. 2025 Aug 26; PMID: 40909733

23. Khan AA, Shekh-Ahmad T, Khalil A, Walker MC, Ali AB. Cannabidiol exerts antiepileptic effects by restoring hippocampal interneuron functions in a temporal lobe epilepsy model. Br J Pharmacol. 2018 Jun;175(11):2097–2115. PMID: 29574880

24. Zhang HXB, Bean BP. Cannabidiol Inhibition of Murine Primary Nociceptors: Tight Binding to Slow Inactivated States of Nav1.8 Channels. J Neurosci. 2021 Jul 28;41(30):6371–6387. PMID: 34131037

25. Ali SR, Liu Z, Nenov MN, Folorunso O, Singh A, Scala F, Chen H, James TF, Alshammari M, Panova-Elektronova NI, White MA, Zhou J, Laezza F. Functional Modulation of Voltage-Gated Sodium Channels by a FGF14-Based Peptidomimetic. ACS Chem Neurosci. 2018 May 16;9(5):976–987. PMID: 29359916

26. Dvorak NM, Wadsworth PA, Aquino-Miranda G, Wang P, Engelke DS, Zhou J, Nguyen N, Singh AK, Aceto G, Haghighijoo Z, Smith II, Goode N, Zhou M, Avchalumov Y, Troendle EP, Tapia CM, Chen H, Powell RT, Baumgartner TJ, Singh J, Koff L, Di Re J, Wadsworth AE, Marosi M, Azar MR, Elias K, Lehmann P, Mármol Contreras YM, Shah P, Gutierrez H, Green TA, Ulmschneider MB, D’Ascenzo M, Stephan C, Cui G, Do Monte FH, Zhou J, Laezza F. Enhanced motivated behavior mediated by pharmacological targeting of the FGF14/Nav1.6 complex in nucleus accumbens neurons. Nat Commun. 2025 Jan 2;16(1):110. PMID: 39747162

27. Zhang HXB, Heckman L, Niday Z, Jo S, Fujita A, Shim J, Pandey R, Al Jandal H, Jayakar S, Barrett LB, Smith J, Woolf CJ, Bean BP. Cannabidiol activates neuronal Kv7 channels. Elife. 2022 Feb 18;11. PMID: 35179483

28. Shen W, Hamilton SE, Nathanson NM, Surmeier DJ. Cholinergic suppression of KCNQ channel currents enhances excitability of striatal medium spiny neurons. J Neurosci. 2005 Aug 10;25(32):7449–58. PMID: 16093396

29. Mikkelsen JD. The KCNQ channel activator retigabine blocks haloperidol-induced c-Fos expression in the striatum of the rat. Neurosci Lett. 2004 May 27;362(3):240–3. PMID: 15158023

30. Lawrence JJ, Saraga F, Churchill JF, Statland JM, Travis KE, Skinner FK, McBain CJ. Somatodendritic Kv7/KCNQ/M channels control interspike interval in hippocampal interneurons. J Neurosci. 2006 Nov 22;26(47):12325–38. PMID: 17122058

31. Filippov AK, Choi RCY, Simon J, Barnard EA, Brown DA. Activation of P2Y1 nucleotide receptors induces inhibition of the M-type K+ current in rat hippocampal pyramidal neurons. J Neurosci. 2006 Sep 6;26(36):9340–8. PMID: 16957090

32. Rivera-Arconada I, Roza C, Lopez-Garcia JA. Enhancing m currents: a way out for neuropathic pain? Front Mol Neurosci. 2009;2:10. PMID: 19680469

33. Docter T, Sorum B, Deshmane R, Doubravsky C, Brohawn SG. Cannabinoid inhibition of mechanosensitive K+ channels. bioRxiv. 2024 Dec 10; PMID: 39713384

34. Talley EM, Solorzano G, Lei Q, Kim D, Bayliss DA. Cns distribution of members of the two-pore-domain (KCNK) potassium channel family. J Neurosci. 2001 Oct 1;21(19):7491–505. PMID: 11567039

35. Heurteaux C, Guy N, Laigle C, Blondeau N, Duprat F, Mazzuca M, Lang-Lazdunski L, Widmann C, Zanzouri M, Romey G, Lazdunski M. TREK-1, a K+ channel involved in neuroprotection and general anesthesia. EMBO J. 2004 Jul 7;23(13):2684–95. PMID: 15175651

36. Monat J, Altieri LG, Enrique N, Sedán D, Andrinolo D, Milesi V, Martín P. Direct Inhibition of BK Channels by Cannabidiol, One of the Principal Therapeutic Cannabinoids Derived from Cannabis sativa. J Nat Prod. 2024 May 24;87(5):1368–1375. PMID: 38708937

37. Martin G, Puig S, Pietrzykowski A, Zadek P, Emery P, Treistman S. Somatic localization of a specific large-conductance calcium-activated potassium channel subtype controls compartmentalized ethanol sensitivity in the nucleus accumbens. J Neurosci. 2004 Jul 21;24(29):6563–72. PMID: 15269268

38. Ji X, Martin GE. BK channels mediate dopamine inhibition of firing in a subpopulation of core nucleus accumbens medium spiny neurons. Brain Res. 2014 Nov 7;1588:1–16. PMID: 25219484

39. Labbaf A, Krauth V, Rychlik N, Narayanan Naik V, Vinnenberg L, Karabatak E, Teasley A, Perissinotti PP, White JA, Meuth SG, van Luijtelaar G, Urbano FJ, Budde T, Zobeiri M. TREK1 Channels Shape Spindle-Like Oscillations, Neuronal Activity, and Short-Term Synaptic Plasticity in Thalamocortical Circuits. J Neurosci. 2025 Nov 5;45(45). PMID: 40998695

40. Bock T, Stuart GJ. The Impact of BK Channels on Cellular Excitability Depends on their Subcellular Location. Front Cell Neurosci. 2016;10:206. PMID: 27630543

41. Wang W, Kiyoshi CM, D. Y, Taylor AT, Sheehan ER, Wu X, Zhou M. TREK-1 Null Impairs Neuronal Excitability, Synaptic Plasticity, and Cognitive Function. Mol Neurobiol. 2020 Mar;57(3):1332–1346. PMID: 31728930

42. Grueter BA, Brasnjo G, Malenka RC. Postsynaptic TRPV1 triggers cell type-specific long-term depression in the nucleus accumbens. Nat Neurosci. 2010 Dec;13(12):1519–25. PMID: 21076424

43. Bisogno T, Hanus L, De Petrocellis L, Tchilibon S, Ponde DE, Brandi I, Moriello AS, Davis JB,Mechoulam R, Di Marzo V. Molecular targets for cannabidiol and its synthetic analogues: effect on vanilloid VR1 receptors and on the cellular uptake and enzymatic hydrolysis of anandamide. Br J Pharmacol. 2001 Oct;134(4):845–52. PMID: 11606325

44. Etemad L, Karimi G, Alavi MS, Roohbakhsh A. Pharmacological effects of cannabidiol by transient receptor potential channels. Life Sci. 2022 Jul 1;300:120582. PMID: 35483477

45. Melas PA, Scherma M, Fratta W, Cifani C, Fadda P. Cannabidiol as a Potential Treatment for Anxiety and Mood Disorders: Molecular Targets and Epigenetic Insights from Preclinical Research. Int J Mol Sci. 2021 Feb 13;22(4). PMID: 33668469

46. Kathmann M, Flau K, Redmer A, Tränkle C, Schlicker E. Cannabidiol is an allosteric modulator at mu- and delta-opioid receptors. Naunyn Schmiedebergs Arch Pharmacol. 2006 Feb;372(5):354– 61. PMID: 16489449

47. Lee JLC, Bertoglio LJ, Guimarães FS, Stevenson CW. Cannabidiol regulation of emotion and emotional memory processing: relevance for treating anxiety-related and substance abuse disorders. Br J Pharmacol. 2017 Oct;174(19):3242–3256. PMID: 28268256

48. Kourrich S, Rothwell PE, Klug JR, Thomas MJ. Cocaine experience controls bidirectional synaptic plasticity in the nucleus accumbens. J Neurosci. 2007 Jul 25;27(30):7921–8. PMID: 17652583

49. Wu X, Shi M, Wei C, Yang M, Liu Y, Liu Z, Zhang X, Ren W. Potentiation of synaptic strength and intrinsic excitability in the nucleus accumbens after 10 days of morphine withdrawal. J Neurosci Res. 2012 Jun;90(6):1270–83. PMID: 22388870

50. Padula AE, Griffin WC, Lopez MF, Nimitvilai S, Cannady R, McGuier NS, Chesler EJ, Miles MF, Williams RW, Randall PK, Woodward JJ, Becker HC, Mulholland PJ. KCNN Genes that Encode Small-Conductance Ca2+-Activated K+ Channels Influence Alcohol and Drug Addiction. Neuropsychopharmacology. 2015 Jul;40(8):1928–39. PMID: 25662840

51. Funke JR, Hwang EK, Wunsch AM, Baker R, Engeln KA, Murray CH, Milovanovic M, Caccamise AJ, Wolf ME. Persistent Neuroadaptations in the Nucleus Accumbens Core Accompany Incubation of Methamphetamine Craving in Male and Female Rats. eNeuro. 2023 Mar;10(3). PMID: 36792361

52. Aceto G, Nardella L, Lazzarino G, Tavazzi B, Bertozzi A, Nanni S, Colussi C, D’Ascenzo M, Grassi C. Acute restraint stress impairs histamine type 2 receptor ability to increase the excitability of medium spiny neurons in the nucleus accumbens. Neurobiol Dis. 2022 Dec;175:105932. PMID: 36427690

53. Zhang XQ, Yu ZP, Ling Y, Zhao QQ, Zhang ZY, Wang ZC, Shen HW. Enduring effects of juvenile social isolation on physiological properties of medium spiny neurons in nucleus accumbens. Psychopharmacology (Berl). 2019 Nov;236(11):3281–3289. PMID: 31197434

54. McDevitt DS, Jonik B, Graziane NM. Morphine Differentially Alters the Synaptic and Intrinsic Properties of D1R- and D2R-Expressing Medium Spiny Neurons in the Nucleus Accumbens. Front Synaptic Neurosci. 2019;11:35. PMID: 31920618

55. Zhang M, Luo Y, Wang J, Sun Y, Xie B, Zhang L, Cong B, Ma C, Wen D. Roles of nucleus accumbens shell small-conductance calcium-activated potassium channels in the conditioned fear freezing. J Psychiatr Res. 2023 Jul;163:180–194. PMID: 37216772

56. Oginsky MF, Maust JD, Corthell JT, Ferrario CR. Enhanced cocaine-induced locomotor sensitization and intrinsic excitability of NAc medium spiny neurons in adult but not in adolescent rats susceptible to diet-induced obesity. Psychopharmacology (Berl). 2016 Mar;233(5):773–84. PMID: 26612617

